# Finding phylogeny-aware and biologically meaningful averages of metagenomic samples: *L*_2_UniFrac

**DOI:** 10.1101/2023.02.02.526854

**Authors:** Wei Wei, Andrew Millward, David Koslicki

## Abstract

Metagenomic samples have high spatiotemporal variability. Hence, it is useful to summarize and characterize the microbial makeup of a given environment in a way that is biologically reasonable and interpretable. The UniFrac metric has been a robust and widely-used metric for measuring the variability between metagenomic samples. We propose that the characterization of metagenomic environments can be achieved by finding the average, a.k.a. the barycenter, among the samples with respect to the UniFrac distance. However, it is possible that such a UniFrac-average includes negative entries, making it no longer a valid representation of a metagenomic community. To overcome this intrinsic issue, we propose a special version of the UniFrac metric, termed *L*_2_UniFrac, which inherits the phylogenetic nature of the traditional UniFrac and with respect to which one can easily compute the average, producing biologically meaningful environment-specific “representative samples”. We demonstrate the usefulness of such representative samples as well as the extended usage of *L*_2_UniFrac in efficient clustering of metagenomic samples, and provide mathematical characterizations and proofs to the desired properties of *L*_2_UniFrac. A prototype implementation is provided at: https://github.com/KoslickiLab/L2-UniFrac.git.

## 1 Introduction

### 1.1 Background and context

Microbes play a vital part in many aspects of our lives, and many diseases and conditions are related to the microbiota in and on our body. Examples of which can include various kinds of cancers [1, 18], as well as in behavioral conditions such as sociability [4] or Autism Spectrum Disorder [5]. Despite holding the key to many of the known problems, metagenomic studies are faced with multiple challenges. One of these difficulties lies in the high variability in metagenomic samples [13, 15]. Since samples for a given environment collected at different sub-environments or at different time points can be vastly different in microbial distribution, it can be difficult to characterize a specific environment by an average or representative distribution.

In this paper, we will present an approach to create such average or representative distributions of a collection of metagenomes by defining a new modification of the classic UniFrac metric [8, 10] which we call the *L*_2_UniFrac. We will look at applications of these representative samples, as well as further providing mathematical characterizations of the *L*_2_UniFrac metric by giving formal mathematical proofs of its properties.

The code used to generate all data and figures in this manuscript can be found in https://github.com/KoslickiLab/L2-UniFrac-Paper.git and a prototype implementation for *L*_2_UniFrac can be found in https://github.com/KoslickiLab/L2-UniFrac.git.

### 1.2 Rationale of approach

In a typical environment-related metagenomics study, clustering is often performed after sample collection and data analysis, the result of which are visualized and analyzed by methods such as Principal Coordinate Analysis (PCoA). It is therefore natural that such clusters can be used as a basis or guide in describing an environment, or in finding a representative distribution that characterizes the environment. Intuitively, it is natural that given a large sample size, the centroid (also called the average or the barycenter) of a clustering of these samples can be used as a representative of the samples in this environment. With such an average, it will be much easier to distinguish the signature microbiomes unique to the environment from random noise that come from microbes rarely in the samples. It will also make the comparison between two environments much easier, simplifying it to the comparison between two sample averages.

Finding the barycenter of a group of distribution is not as trivial as it seems. For the purpose of scalability and practicality, the method of computing such a barycenter should be relatively easy and the result easily interpretable biologically. The key to both of these properties lies in the choice of distance metric. A clustering process like this requires a distance metric such that the dissimilarity between two microbial distributions, a property also known as the beta-diversity, can be measured. There are multiple such beta-diversity metrics. In this paper, we will focus on the discussion of finding the barycenter with respect to the UniFrac metric.

There are two main reasons why the UniFrac metric was chosen among the others. The first is due to its robustness. Being one of the most widely-used phylogenetic metrics, the UniFrac metric has demonstrated its usefulness in many areas ranging across clinical studies [7], environmental studies [14], and forensic science [16]. Compared to non-phylogenetic metrics such as the Jaccard index and the Bray-Curtis dissimilarity index, the UniFrac metric takes into account the phylogenetic relationships among the organisms, giving it more biological context and hence more interpretability. In real applications, the UniFrac metric has also demonstrated its superiority over other methods, such as in sensitivity in the identification of enterotypes, which are subtypes of the same environment [7], making it an ideal candidate. The second reason is due to the unique and interesting mathematical characterization of UniFrac, which makes this biologically-motivated problem mathematically interesting at the same time, and leaves rooms for mathematical generalization that could potentially be useful for other related problems yet to be explored.

We begin by first understanding this mathematical characterization and how it is related to our goal of finding the barycenter with respect to the UniFrac distance. About seven years after the development of the original UniFrac in 2005 [8], it was shown by Evans et al. that the UniFrac metric is closely related to a mathematical problem that had been studied for centuries: the optimal transport problem [2]. The optimal transport problem uses a distance known as the Wasserstein distance, or the Earth Mover ‘s distance. Evans showed that the UniFrac distance is equivalent to the 1-Wasserstein distance over a phylogenetic tree. Under this formulation, instead of the original computation of UniFrac as a fraction of branch lengths, it can be equivalently computed by representing metagenomic samples by probability distributions and “aggregating” the distributions up a phylogenetic tree through matrix multiplication, obtaining the aggregated vectors in what we will call the “*L*_1_UniFrac space”, followed by taking the *L*_1_ norm between two such aggregated vectors. This alternative characterization of UniFrac, termed the “*L*_1_UniFrac” by Koslicki and McClelland [9], has several benefits. First, it allows an alternative representation of a metagenomic sample as an aggregated vector in the *L*_1_UniFrac space, in which the distance between two vectors can be easily computed by taking the *L*_1_-norm. Also, this representation of data points in the *L*_1_UniFrac space leaves room for computation of the *L*_1_-mean, or median, among a group of samples, giving rise to an easy-to-compute centroid, which can be viewed as the average of the samples in the *L*_1_UniFrac space. This idea of computing the “*L*_1_-mean UniFrac” was first mathematically conceptualized by Koslicki and McClelland [9], who saw it as an opportunity to find the representative sample of an environment with respect to the UniFrac metric.

However, the “*L*_1_-mean sample” computed using this method does have a detrimental issue. We want the mean/average to have a readily interpretable biological meaning of representing an average sample in the environment, and not simply as an aggregated vector in the *L*_1_UniFrac space. The former requires it being a distribution vector with each entry being non-negative and all the entries sum up to 1, such that each entry represents the relative abundance of an organism in the sample. This desired property cannot be guaranteed when the mean vector in the *L*_1_UniFrac space is projected back to the distribution space. Both experiments using real world data and mathematical deductions have shown that the resulting projected vector could contain negative entries [12]. This makes the result biologically meaningless. Our proposed solution is an alternative characterization of UniFrac using the *L*_2_ norm instead of *L*_1_. Without changing the nature of being a solution to an optimal transport problem, the so-termed *L*_2_UniFrac retains the original spirit of UniFrac of being a phylogeny-aware dissimilarity metric and offers comparable robustness, while circumventing the issue of having negative values in *L*_1_UniFrac mean.

## 2 Methods

Herein, we describe the *L*_2_UniFrac metric and how to use it to take averages. Full details and proofs are provided in the Supplementary material.

### 2.1 *L*_1_ **and** *L*_2_ **Unifrac**

Given a phylogenetic tree *T* with *N* ordered nodes, a metagenomic sample can be represented as a probability distribution *P* = (*P*_1_, *P*_2_, …, *P*_*N*_) with entries in the same order as the nodes of *T*, such that *P*_*i*_ represents the relative abundance of organism/taxa *i* in sample *P*.

Define

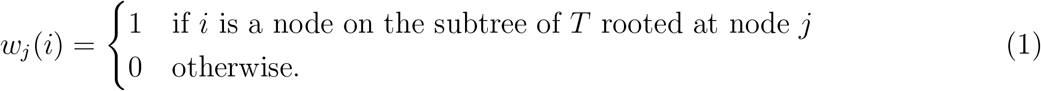

Let *W* be an *N × N* matrix with row *j* being *w*_*j*_ scaled by length of the branch connecting node *j* and its ancestor. The matrix 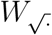. is defined similarly, but with the rows scaled by the square root of the branch length.

We can now recall the definition of the UniFrac metric (see, for example, and of [9, 11, 2]):

**Definition 1** (UniFrac). For two metagenomic samples represented by the probability distributions *P* and *Q*, and with *W* as defined above for some fixed phylogenetic tree, and for || *·* ||_1_ the standard *L*_1_ norm,

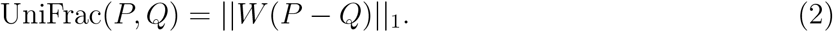

By simply changing the norm from the *L*_1_ to *L*_2_ norm, we obtain:

**Definition 2** (*L*_2_UniFrac). For two metagenomic samples represented by the probability distributions *P* and *Q*, and with 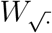 as defined above for some fixed phylogenetic tree, and for || *·* ||_2_ the standard *L*_2_ norm,

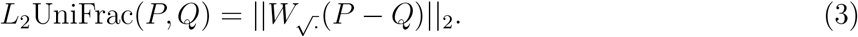

In the Supplementary material, we demonstrate that one can take meaningful averages with respect to the *L*_2_UniFrac metric, but not with the traditional (*L*_1_) UniFrac.

In practice though, we do not need to form the large matrix 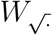., but rather proceed in a post-order aggregation to implement the matrix multiplications 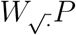. and 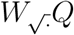. in equation (3) in Definition 2: see Algorithm 1. After applying Algorithm 1 to compute 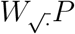 *P* and 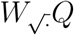 *Q*, the

#### Algorithm 1

*L*_2_-aggregate: an algorithm to obtain the *L*_2_-aggregated vector given a probability vector *P*, resulting in a vector in *L*_2_UniFrac space.

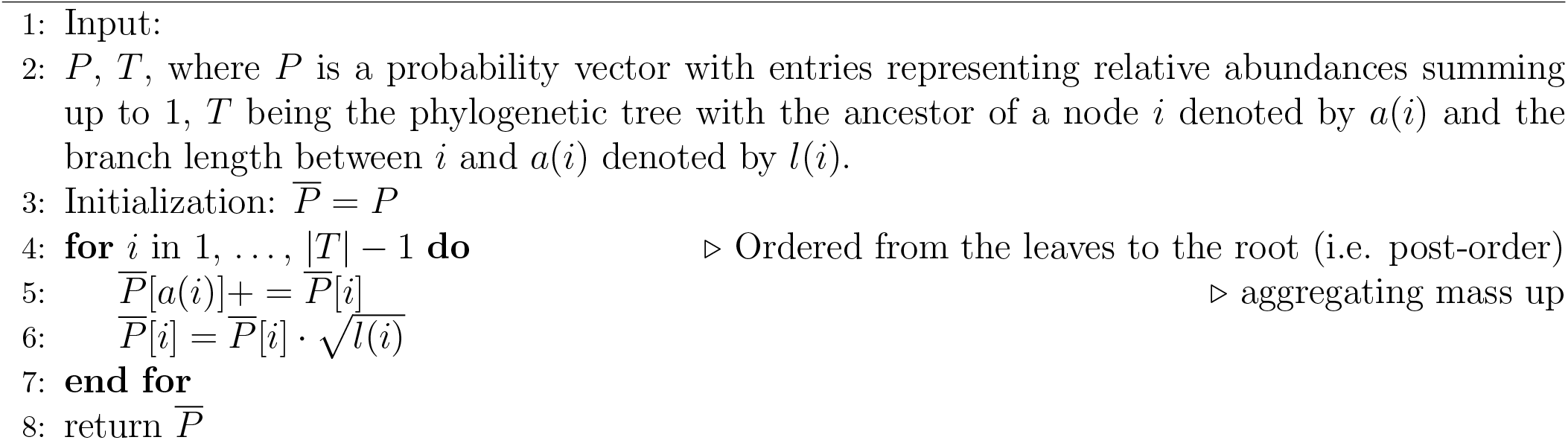

*L*_2_UniFrac can then be computed by taking the simple *L*_2_-norm of the difference of the resulting vectors.

Later, we will need to also compute the inverse of this operation (i.e. compute 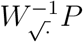 and 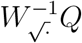), the algorithm for which is similar to the above; see Algorithm 2.

#### Algorithm 2

Inverse-aggregate: an algorithm that reverse *L*_2_-aggregate to obtain a probability vector in the original space, given an aggregated vector in the *L*_2_UniFrac space.

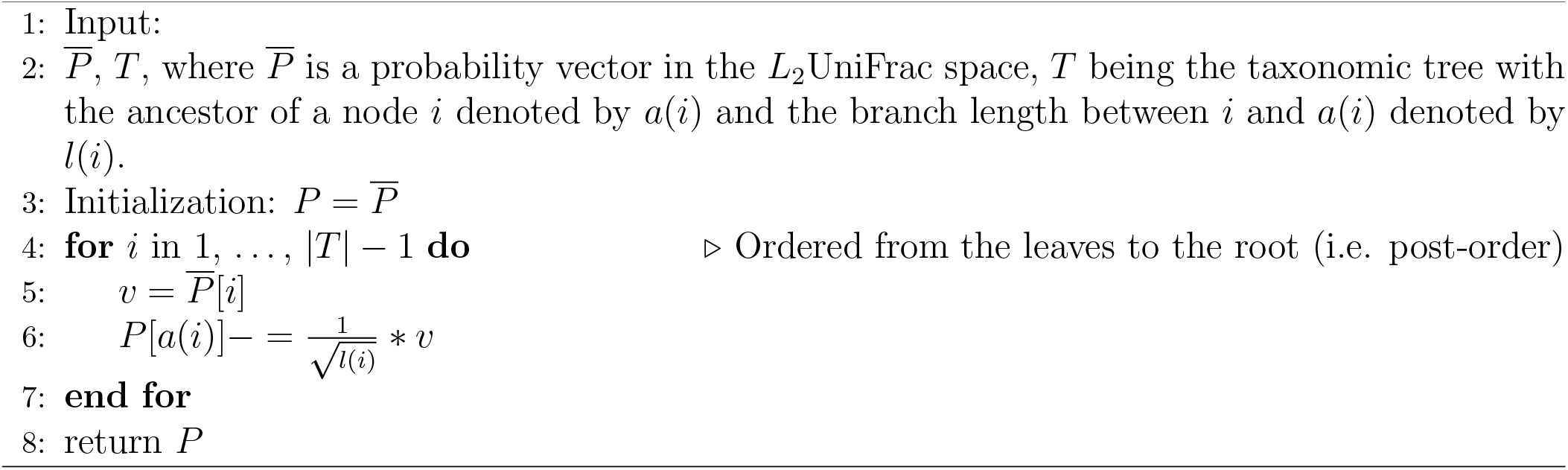

### 2.2 Computing *L*_2_UniFrac averages

In the Supplementary material, we demonstrate that one cannot form biologically meaningful (*L*_1_) UniFrac averages of a collection of metagenomic samples represented by a collection of probability vectors. In short, this is due to the probability simplex not being closed under the operation of taking medians. However, the probability simplex is closed under means (See Supplementary material section S1.4). As such, we describe taking averages with respect to the *L*_2_UniFrac metric only.

The definition of an average (or more precisely, a barycenter) with respect to the *L*_2_UniFrac metric is as follows:

**Definition 3** (*L*_2_UniFrac barycenter). Given a collection of probability distributions 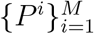 representing metagenomic samples, each of which is given by an *N* -length vector indexed by the nodes in a fixed phylogenetic tree *T*, the average, or barycenter, of the set 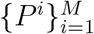 is given by *P*^*∗*^

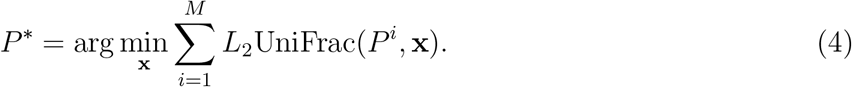

Thus, in practice, to compute a “representative sample” of a collection of vectors 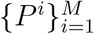, w proceed as follows: First, form 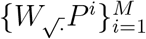 by repeat application of Algorithm 1. Then, form the *L*_2_UniFrac barycenter *P*^*∗*^ by taking the component-wise mean of the collection 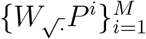. Finally, use Algorithm 2 to compute 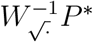, the result of which will be a probability distribution representing the *L*_2_UniFrac average of the collection 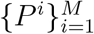. Most interestingly, we show in the Supplementary material, Claim 4 that the result of this process is as simple as taking the componentwise mean of the collection of vectors 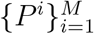.

For two collections of vectors (representing, say, two different environments) 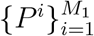 and 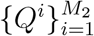. the “representative sample” vectors 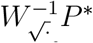 and 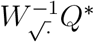 can be used as a proxy of their respective collections. Thus, instead of needing to form all pairwise *L*_2_UniFrac distance calculations as:

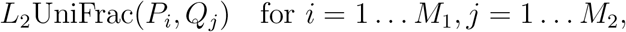

and averaging over the environments, we can much more efficiently do the single computation:

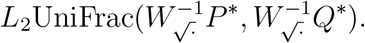

This amounts to applying Algorithm 1 to the component-wise averages of 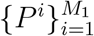 and 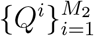 then taking the *L*_2_-norm of the difference between these two vectors.

In the following, we will refer to “*L*_*p*_UniFrac.” By this we mean the following: for a sample distribution *P* (living in the probability simplex), left-multiplying by the aforementioned *W* (modified depending on the *L*_*p*_ space under consideration), we obtain a vector *WP* living in “*L*_*p*_UniFrac space.”

## 3 Results

### 3.1 Clustering in *L*_2_UniFrac space

We hypothesize that phylogeny-aware clustering of samples, which is a common procedure in metagenomic analyses, can be done much faster in the *L*_*p*_UniFrac space than in the distribution space. The reason is as follows: The traditional way of UniFrac-based clustering (in the distribution space) requires the pairwise UniFrac distances be computed prior to applying clustering methods. The computation of UniFrac is not trivial. Pairwise comparison aggravates the cost of computationM by an exponential factor as sample size increases. On the other hand, when samples are represented as aggregated vectors in the *L*_*p*_UniFrac space, the UniFrac distance can be readily computed by simply computing the *L*_*p*_-norm. The bulk of the computation lies only in transforming *P* to *WP* through matrix multiplication, which is linear with respect to sample size.

We tested this hypothesis using *L*_2_UniFrac as a representative of the *L*_*p*_UniFrac general case. To demonstrate the improvement on clustering speed, we randomly selected 1,000 samples out of the 6,067 samples of 16S data obtained from Qiita [3] (study ID 1928), consisting of samples collected from four body sites: skin, saliva, vagina, and feces. Out of these samples, we randomly sampled with sample size ranging from 50 to 800 in step of 50. Each of these samples were clustered using two methods: the conventional matrix-based method, in which a pairwise UniFrac distance matrix is computed using the EMDUniFrac [10] algorithm, followed by *k*-medoids clustering, as well as our proposed method, in which samples were first aggregated to the *L*_2_UniFrac space, followed by *k*-means clustering on aggregated vectors. We further computed the Fowlkes-Mallows score for each of the instances as a measure of the clustering quality. The comparisons of both the time cost and the clustering quality are shown in Figure 1.

**Figure 1:**
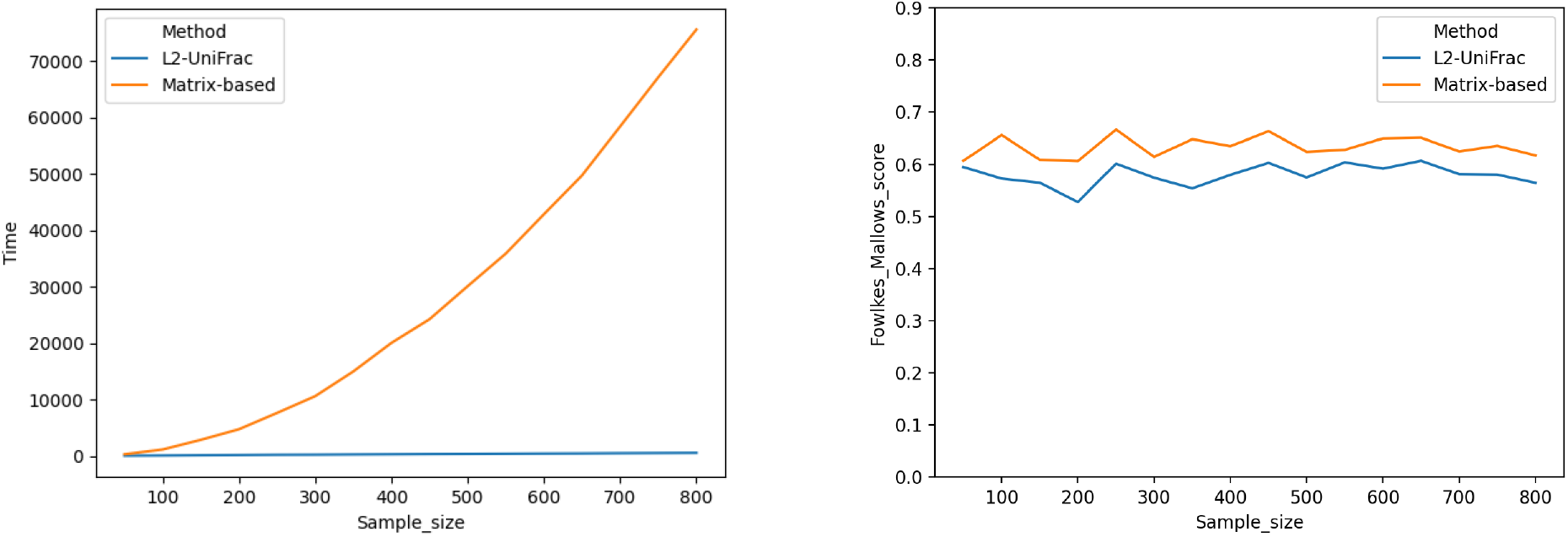
Comparison between the performance of the traditional matrix-based clustering, where pairwise UniFrac distance matrix was computed first and used as the basis for clustering, versus *L*_2_-clustering, where data points were directly clustered in the *L*_2_UniFrac space using *L*_2_ norm as a distance metric. Left: running time. Right: clustering quality measured using Fowlkes-Mallows score, with a higher score indicating a better clustering.

As expected, for the traditional method, clustering time increased exponentially as sample size increases. According to the figure, for as few as 300 samples, the traditional matrix-based method would require approximately 3 hours to complete, whereas clustering in *L*_2_UniFrac space took only a few minutes. This improvement on speed does come with a trade-off in clustering quality, as suggested by the figure on the right. However, the difference in clustering score is only around 0.1 on a scale of 0 to 1 and remained fairly constant in the experiments performed. In the case of large data size where the exact accuracy is not of top priority, the significant improvement in speed would deem clustering on *L*_2_UniFrac space much more practical than the traditional ‘all pairwise UniFrac” approach in real life scenarios.

### 3.2 Environment fingerprinting and classification

#### 3.2.1 Finding the representative sample

In this section, we illustrate the process of finding the average sample using *L*_2_UniFrac specific algorithms.

The illustration was performed using data with study ID 714 from `Qiita` [3], consisting of 528 samples from different environments. We first performed Algorithm 1 on all samples to obtain the aggregated vectors in the *L*_2_UniFrac space. We then took the component-wise *L*_2_ mean of these vectors. This mean vector was projected back to the distribution space using Algorithm 2, obtaining an average sample of the original samples. Principal Coordinate Analysis (PCoA) plots were used to show the relative relationship among these samples. Out of the five environments, we removed two that had too few samples and singled out each environment together with its representative data point to better observe their relationship for the remaining three. The results are shown in Figure 2. From the figure, a representative sample corresponds roughly to the centroid of the cluster consisting of the samples belonging to that environment. It turns out that this average sample can be equivalently computed by simply computing the component-wise mean of the original distributions, as shown in the Supplementary material Section S1.4.2. This further simplifies and speeds up the computation and is a unique advantage of *L*_2_UniFrac that cannot be achieved using *L*_1_UniFrac (see Supplementary material Section S1.4.1), making *L*_2_UniFrac more than simply an alternative version of UniFrac but instead a novel metric having its own unique applications and properties.

**Figure 2:**
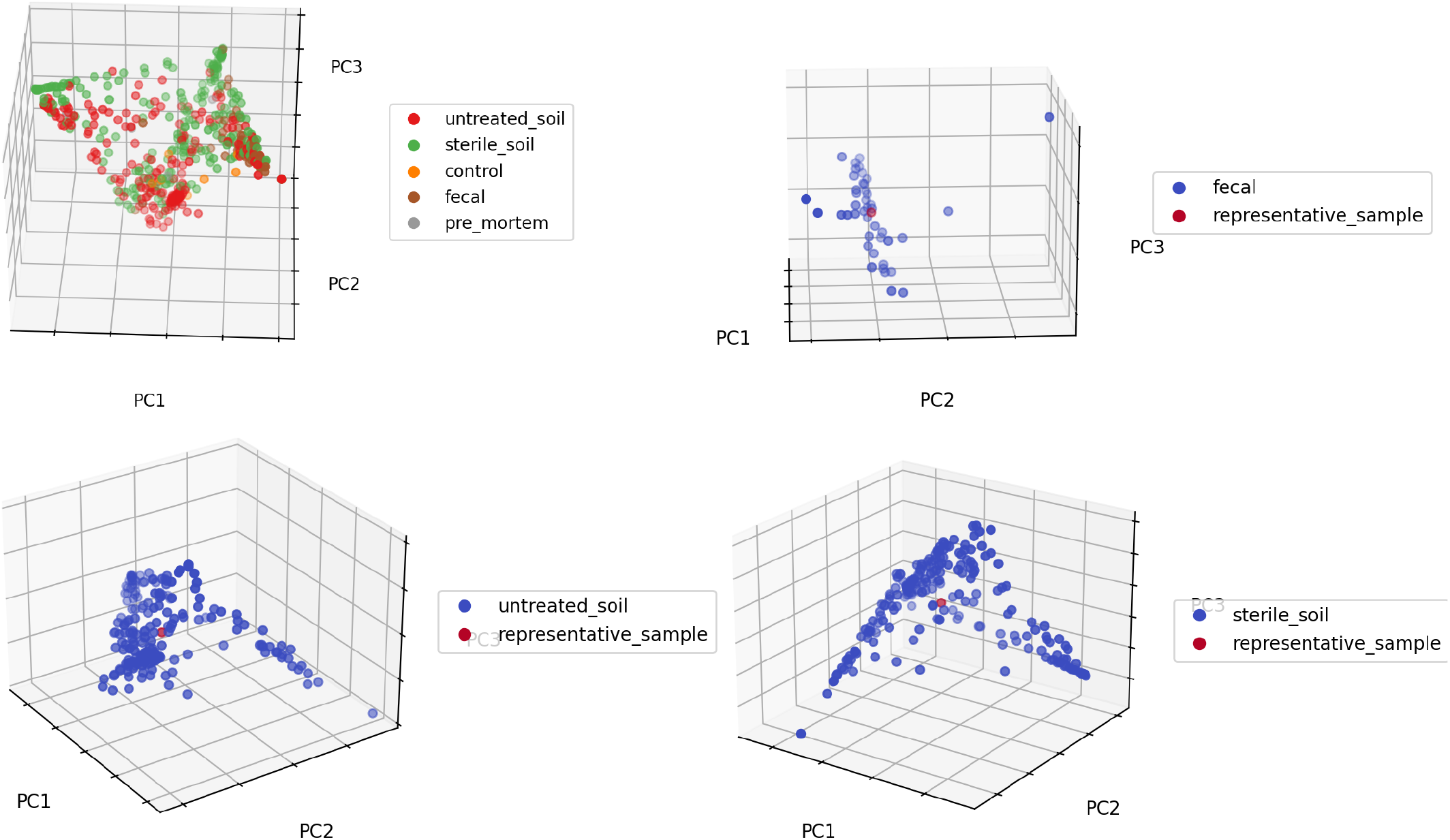
PCoA plots showing the representative samples with respect to all the samples from the respective environments.

#### 3.2.2 Classification

Up to this point, we have demonstrated the process of obtaining the average sample amongst a pool of samples from the same environment. Ideally, this representative sample should be specific to the environment such that there is sufficient degree of differences between the representative samples of two different environments to significantly distinguish one from another. In this section, we test this hypothesis using a classification test using the same data set from Section 3.1 consisting of 16S samples from four body sites. Typically, when a group of unknown samples are given, clustering is first performed and classes are assigned based on the clustering result, such as based on the minimum distance to the centroid of the clusters, or by majority vote from a certain number of closest points. This type of methods can be time consuming and the unsupervised nature of this method also does not guarantee a direct correspondence between the segregation of the clusters and that of the actual traits of interest. The alternative we propose here is to first find the representative samples corresponding to each environment using a large size of known sample. For any subsequent new sample, its class can be assigned based on similarity with the representative samples. The accuracy of such assignments can therefore be used to gauge the specificity of such representative samples.

To this end, we partitioned the samples of each body site into 80 percent of training samples and 20 percent of testing samples. As representatives of current clustering-based methods, we considered *k*-medoids and *k*-means as clustering methods. For each of these clustering methods, we first performed clustering, using the pre-computed pairwise *L*_2_UniFrac distance matrix for *k*-medoids and probability distribution vectors for *k*-means, setting the number of clusters equal to the number of body sites present. With 80 percent of the training data, the label for each cluster was assigned by majority vote. For each testing sample, the labels assigned by clustering was compared against its true label. For our *L*_2_UniFrac based method, a representative sample was first computed using the training data. We then computed the *L*_2_UniFrac distance between each of the testing samples and each of the representative samples and assign the test sample to the body site of which the *L*_2_UniFrac between the representative sample produced the minimum *L*_2_UniFrac distance. We used different scoring metrics from the `sci-kit learn` python package to evaluate the performance of all three classification methods. Figure 3 shows one such score, accuracy, used. The performance evaluated using other scores is shown in Figure (Supplementary Figure S1).

**Figure 3:**
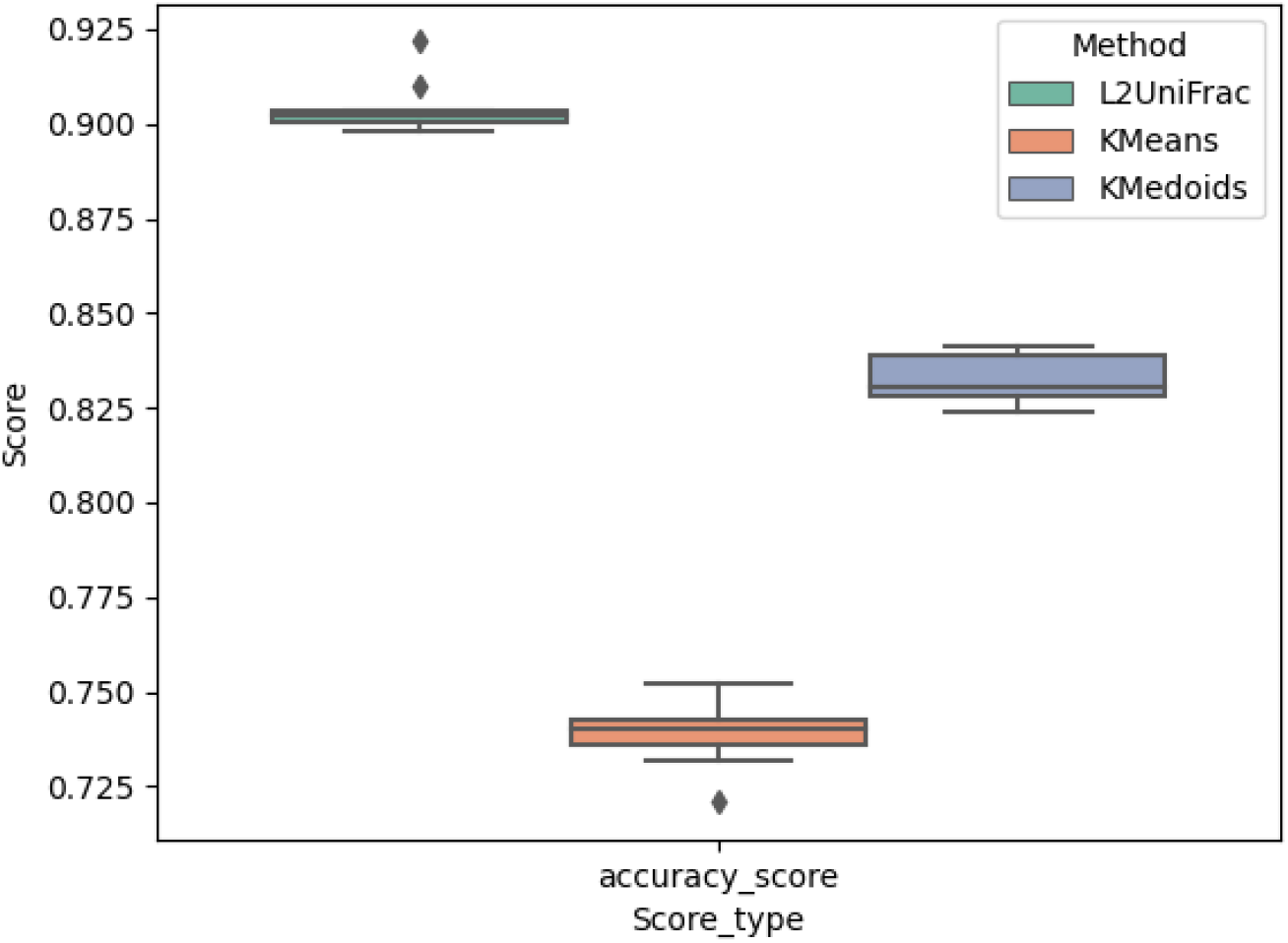
The accuracy scores of three classification methods. The accuracy score is defined as the proportion of correct classifications over approximately 1210 classifications.

The result shows that our method out-performed the other two methods in most cases. The poor performance of *k*-medoids and *k*-means can be partially attributed to the fact that the labels assigned by majority vote after clustering were not consistent with the original classes, though the number of clusters were set to be equivalent to the true number of classes. For instance, in some cases, two clusters were assigned the same label by majority vote, while some of the original classes were missing in this process. This significantly affected the results.

Of course, this comparison is between two different kinds of approaches: supervised and unsupervised. In the case of the *L*_2_UniFrac method, the exact classes were known and the representative samples were created from samples from a known environment. In the case of *k*-means and *k*-medoids, however, the clustering was unsupervised, significantly increasing the proportion of false positives and false negatives. On the other hand, this in turn shows the very advantage of our method of having the potential to convert a traditionally more unsupervised method into one with a more supervised nature by creating a “reference point” for each environment.

#### 3.2.3 Identifying differentially abundant organisms

An additional motivation of finding the average sample that characterizes an environment lies in the hope of identifying signature microbiomes that distinguish one environment significantly from another. Traditionally, in order to identify the most differentially abundant organisms, the abundances of each organism are to be compared across all samples. The results can then be visualized commonly using a box and whisker plot showing the average abundances of the organisms in each sampling environment, based on which a conclusion can be drawn on if the differences are significant, with the aid of statistical methods. We propose the alternative method based on the “flow” between average samples obtained through *L*_2_UniFrac, giving rise to a phylogeny-aware comparison between sample groups.

The formulation of UniFrac as an optimum transport problem allows the differences in two samples to be represented as the “flow” between the two distributions. More specifically, for two samples (or averages) *P* and *Q*, the vector 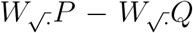 represents the flow (or flux) across0020 edges in the phylogenetic tree when moving abundance from *P* to overlap that of *Q*. This vector 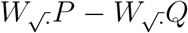 is called the differential abundance vector (see [10] for the *L*_1_ analog). The larger the absolute value of the flow between two organisms at a certain node in the phylogentic tree, the greater the difference is between the two organisms at this node/taxa. As such, we propose to obtain first the average samples from each environment and then compute the flow between pairwise average samples for the detection of differentially abundant taxa.

To test the feasibility of our method, we used the PRJEB6070 (BioProject 266076) study downloaded from HumanMetagenomeDB [6], consisting of 1261 gut samples grouped by three conditions: colorectal cancer, adenoma, and control. Unlike the previous experiments using 16S rRNA data, this dataset consists of whole genome shotgun (WGS) data. Using the principle adopted by the WGSUniFrac method [17], the average abundance with respect to each taxonomic ID is computed for each condition, giving rise to three representative samples. Using the taxonomic tree in place of the phylogenetic tree with the reciprocal-based assignment of branch lengths as suggested by the WGSUniFrac method [17], the pairwise flow among the three representative samples were computed and illustrated in Figure 4.

**Figure 4:**
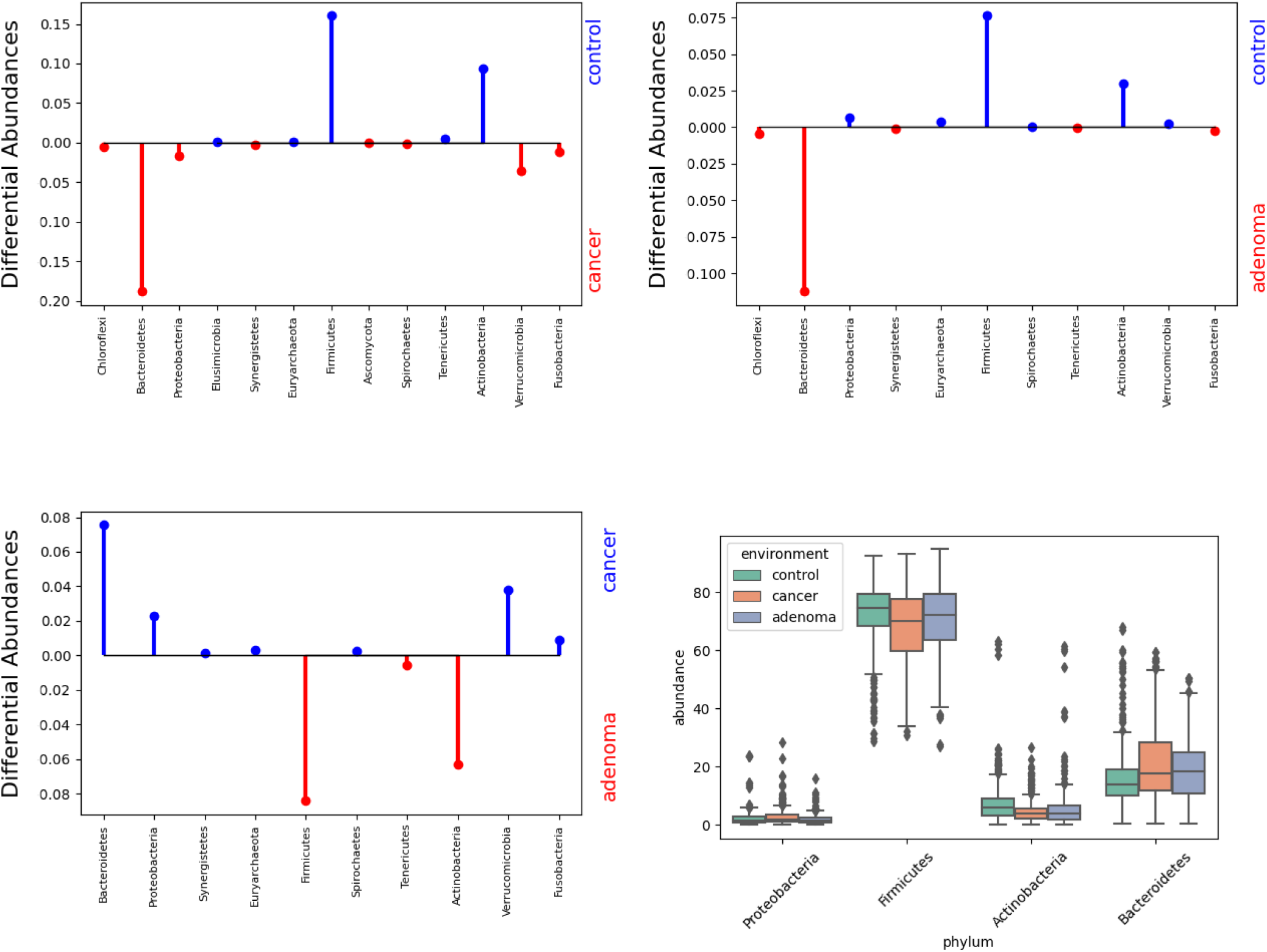
Top two and bottom left: top differentially abundant phyla between each of the condition pairs. The values are measured in terms of the *L*_2_UniFrac flow. Bottom right: a traditional box plot of taxa relative abundances showing the spread of selected phyla.

Based on the differential abundance plots, we identified Bacteroidetes, Firmicutes, Actinobacteria, and Proteobacteria to be some of the phyla with the highest degree of distinguishability across the three conditions. The differential abundances plots show that compared to control, cancer and adenoma conditions tend to have a higher abundance of Bacteroidetes while having a lower abundance in Firmicutes and Actinobacteria. These agree with the box plot in the same Figure 4. Interestingly, these phyla had been observed in a previous study [18], supporting the sensitivity of our method. However, though the same phyla were identified to be correlated with the conditions, the trend of correlation does not seem to agree, with our data showing an increased abundance in Bacteroidetes and decreased abundances in Firmicutes and Actinobacteria in adenoma and cancer conditions compared to the control group. The study carried out by Yachida et al. in [18] demonstrated elevation in all of the above-mentioned phyla. This is likely due to different data sets being compared. Perhaps more studies were to be conducted to reach a decisive conclusion. One thing we do agree, though, is the fact that the observed patterns seems to be progressive, with adenoma falling in between control and cancer, making these phyla strong candidates as indicators of the development of colorectal cancer.

## 3.3 Discussion

In this paper, we proposed *L*_2_UniFrac as an alternative of the traditional *L*_1_UniFrac. This *L*_2_UniFrac preserves the robustness of *L*_1_UniFrac as a phylogenetic metric for beta-diversity and allows one to compute the average distribution with respect to *L*_2_UniFrac metric, which is not possible with the original version of UniFrac. Furthermore, we further explored the properties of the *L*_2_UniFrac space, in which lie the aggregated vectors obtained by aggregating the distributions up the phylogenetic tree. In this space, clustering of metagenomic samples can be performed with much higher efficiency, circumventing the computation of the pairwise distance matrix, with negligible sacrifice of clustering quality. Given the invertibility of the aggregation process, as shown in Section S1.2, the *L*_2_-mean of the vectors in the *L*_2_UniFrac space can be taken and projected back to the distribution space, giving rise to an average sample with respect to *L*_2_UniFrac. We further showed that this projected mean is also a probability distribution, and is actually none other than the component-wise mean of the metagenomic distributions. This property gives the component-wise average of metagenomic distributions a phylogeny-aware biological interpretation as the *L*_2_ norm of their differences is exactly the *L*_2_UniFrac value.

We also demonstrated some of the potential usage of such average samples through experimentation. Most significantly, such average samples can serve as fingerprints for specific environments, allowing a biologist to better characterize the signature microbes belonging to a specific environment. Our results in Section 3.2 show that despite the between-sample variability even within the same environment, the average sample is sufficient to distinguish one environment from another to a fair degree. The average sample obtained using this method is stable to the environment in the sense that given the same environment and sufficiently large sample size, the variance between different average samples obtained using different sample pools or at different time points can be minimized. The reason is due to the nice property of its equivalence with the component-wise *L*_2_-mean, making this process equivalent to taking a sample mean. By the Central Limit Theorem, when the sample size is sufficiently large, the sample means will be approximately normally distributed, with a standard deviation inversely proportional to the number of times samples are taken. Of course, the difficulty in defining an environment still remains, such as in the case of disease development, where different stages of disease are often artificially defined based on physiological differences. However, it can be envisioned that by pooling samples from adjacent stages, the average sample of intermediate stages can be computed. Coupled with the method to compute the flow between two distributions as demonstrated in Section 3.2.3, phylogeny-aware differential abundance profiles can be obtained, with the potential to provide a trajectory on the dynamic changes in metagenomic diversity with respect to the development of the disease to user-defined resolution.

## Supporting information

Mathematical Supplementary Materials

## 4 Acknowledgment

This work was supported by the NIH grant 1R01GM146462-01.

