## Supplementary material for "Finding phylogeny-aware and biologically meaningful averages of metagenomic samples: *L*_2_UniFrac": Mathematical Supplementary Materials

### S1 Mathematical descriptions and proofs

#### S1.1 Definitions and terminology

We recall the following definitions of $L_1$ and $L_2$ UniFrac from the Methods section of the main manuscript.

##### S1.1.1 $L_1$UniFrac

In the original description of UniFrac distance as 1-Wasserstein distance, each sample is represented as a probability vector in $\mathbb{R}^N$ where $N$ is the number of nodes on the phylogenetic tree. Given such two vectors $P$ and $Q$, the UniFrac distance between them can be computed by by computing

$$\text{UniFrac}(P, Q) = ||W(P - Q)||_{L_1}$$

where $W$ is an $N \times N$ matrix with the $i$-th row being the indicator function describing the subtree rooted at node $i$ scaled by length of the branch connecting node $i$ and its ancestor (see section S1.2 below).

##### S1.1.2 $L_2$UniFrac

The $L_2$UniFrac can be expressed in a similar manner, with the $L_1$ norm replaced by $L_2$ norm and $W$ scaled by the square-root of branch lengths instead. The scaling of branch lengths by taking square-root is motivated by the biological meaning that would result from this expression as noted by Evans and Matsen [2, 6]. Since the branch lengths are simply a measure of phylogenetic proximity, taking the square-root will not change the underlying principle of this computation. We denote this expression by

$$L_2\text{UniFrac}(P, Q) = ||W_{\sqrt{\cdot}}(P - Q)||_{L_2}. \tag{1}$$

For any distribution $P$, we will refer to the metric spaces where $WP$ and $W_{\sqrt{\cdot}}P$ lie in the $L_1$UniFrac space and the $L_2$UniFrac space respectively.

#### S1.2 Invertibility of $W$ and $W_{\sqrt{\cdot}}$

**Claim 1.** *For any given tree with non-zero branch lengths, the corresponding $W$ or $W_{\sqrt{\cdot}}$ constructed as above is invertible.*

*Proof.* We proof this claim by showing that $W$ and $W_{\sqrt{\cdot}}$ are lower triangular matrices with non-zero diagonal entries, indicating that they are full-rank matrices and are thus invertible.

To construct $W$ or $W_{\sqrt{\cdot}}$ given a tree $T$ with $N$ nodes, we first define an indicator function as follows:

$$w_j(i) = \begin{cases} 1 & \text{if } i \text{ is a node on the subtree of } T \text{ rooted at node } j \\ 0 & \text{otherwise.} \end{cases} \tag{2}$$

Let $w_j$ be a row vector with entry $i$ being $w_j(i)$ for $1 \leq i \leq n$. Then $w_j$ will have 1 on position $j$ as well as all descendants of $j$, and 0 everywhere else. In particular, $w_j(i) = 0 \; \forall i \geq j$. Let $W'$ be a matrix with row $j$ being $w_j$ for $1 \leq j \leq n$. Then it can be observed that $W'$ is an $N \times N$ lower-triangular matrix with 1 on the diagonal, and is thus invertible. Now $W$ (or $W_{\sqrt{\cdot}}$) is constructed by scaling $W'$ row by row by a positive factor (some non-zero function of the length of the corresponding branch, be it the branch length itself or the square-root of it). The resulting matrix is still a lower-triangular matrix with non-zero diagonal entries, and is hence invertible. □

In the next section, we present the algorithms that compute $W_{\sqrt{\cdot}}$ and its inverse respectively, and the utilities of these operations.

#### S1.3 Motivation and computations of $W_{\sqrt{\cdot}}$ and $W_{\sqrt{\cdot}}^{-1}$

The first motivation for the computation of $W_{\sqrt{\cdot}}$ is for the purpose of clustering in $L_2$UniFrac space, as demonstrated in Section 3.1. As the definition (1) of $L_2$UniFrac shows, the $L_2$UniFrac distance between two distribution vectors $P$ and $Q$ can be equivalently expressed as $||W_{\sqrt{\cdot}}P - W_{\sqrt{\cdot}}Q||_{L_2}$, which is simply the $L_2$ distance between two $W_{\sqrt{\cdot}}$-transformed vectors. Instead of constructing the actual matrix $W_{\sqrt{\cdot}}$ and perform matrix multiplication, which is both time-consuming and space-consuming, Algorithm 1 performs this exact process in a bottom-up fashion with respect to tree $T$, resulting in a linear-time and linear-space performance. As this algorithm can be perceived as aggregating the masses from leaves up the tree $T$, resulting in an 'aggregated vector' in $L_2$UniFrac space, we term this algorithm '$L_2$-aggregate' to illustrate this property. The correctness of a very similar algorithm, but the $L_1$UniFrac case, is shown by Wei and Koslicki in [13], which can be directly adopted to demonstrate the correctness of Algorithm 1 by substituting $l(i)$ with $\sqrt{l(i)}$.

**Algorithm 1** $L_2$-aggregate: an algorithm to obtain the $L_2$-aggregated vector given a probability vector $P$

Input:
$P$, $T$, where $P$ is a probability vector with entries representing relative abundances summing up to 1, $T$ being the phylogenetic tree with the ancestor of a node $i$ denoted by $a(i)$ and the branch length between $i$ and $a(i)$ denoted by $l(i)$.
Initialization: $\overline{P} = P$
**for** $i$ in 1, . . . , $|T| - 1$ **do** ▷ Ordered from the leaves to the root
  $\overline{P}[a(i)] += \overline{P}[i]$ ▷ aggregating mass up
  $\overline{P}[i] = \overline{P}[i] \cdot \sqrt{l(i)}$
**end for**
return $\overline{P}$

Similarly, we have Algorithm 2 that computes $W_{\sqrt{\cdot}}^{-1}$, which we termed 'inverse $L_2$-aggregate' as it reverses the effect of Algorithm 1 to obtain the original vector $P$ from its aggregated counterpart $\overline{P}$.

**Algorithm 2** Inverse-aggregate: an algorithm that reverse $L_2$-aggregate to obtain a probability vector in the original space, given an aggregated vector in the $L_2$UniFrac space.

Input:
$\overline{P}$, $T$, where $\overline{P}$ is a probability vector in the $L_2$UniFrac space, $T$ being the taxonomic tree with the ancestor of a node $i$ denoted by $a(i)$ and the branch length between $i$ and $a(i)$ denoted by $l(i)$.
Initialization: $P = \overline{P}$
**for** $i$ in 1, . . . , $|T| - 1$ **do** ▷ Ordered from the leaves to the root
  $v = \overline{P}[i]$
  $P[a(i)] -= \frac{1}{\sqrt{l(i)}} * v$
**end for**
return $P$

It is noteworthy that the existence of Algorithm 2 is not merely for mathematical completeness, but has the actual application of converting an aggregated vector $\overline{P}$ in the $L_2$UniFrac space to the corresponding distribution vector $P$ in the distribution space, even if $P$ is not a naturally-existing distribution by itself.

For instance, $\overline{P}$ can be a vector that is obtained by applying Algorithm 1 to a group of samples belonging to the same environment and taking the average of these $L_2$-aggregated vectors. Algorithm 2 allows one to obtain the corresponding distribution $P$, which can be seen as an 'average' distribution representing the environment.

#### S1.4 Obtaining the average sample in $L_1$ and $L_2$ UniFrac space

We first give a formal definition to average distribution in a metric space.

**Definition 1** (Barycenter)**.** Let $X = \left(\mathbb{R}^N, d\right)$ be a metric space, the barycenter $x^*$, or the average distribution of $n$ probability distributions $x^1$, $x^2$ ..., $x^n$ in $X$ is defined by:

$$x^* = \arg\min_x \sum_{i=1}^{n} d(x^i, x). \tag{3}$$

**Claim 2.** *The barycenter of a finite number of vectors in $L_1$ space is equivalent to the median of the vectors.*

*Proof.* Let $x^1$, $x^2$, ..., $x^n \in (\mathbb{R}^N, |.|)$ be $n$ vectors in $L_1$ space, where $|.|$ is the $L_1$-norm. Then the barycenter $x^*$ satisfies

$$x^* = \arg\min_x \sum_{i=1}^{n} |x - x^i| \tag{4}$$

$$\Rightarrow x_j^* = \arg\min_{x_j} \sum_{i=1}^{n} |x_j - x_j^i| \quad \forall 1 \leq j \leq N. \tag{5}$$

$$\tag{6}$$

Suppose $x_j^*$ is not the median of $x_j^1, x_j^2, ..., x_j^n$. Then, assume without loss of generality that $p$ out of $n$ $x_j$'s are smaller than $x_j^*$ and $q$ out of $n$ $x_j$'s are greater than or equal to $x_j^*$, such that $p + q = n$ and $p < q$. Let $\overline{x}_j$ be the smallest number greater than $x^*$ and let $d = |\overline{x}_j - x_j^*|$. Then

$$\sum_{i=1}^{n} |\overline{x}_j - x_j^i| = \sum_{i=1}^{n} |x_j^* - x_j^i| + pd - qd < \sum_{i=1}^{n} |x_j^* - x_j^i| \tag{7}$$

which is a contradiction. □

##### S1.4.1 The insufficiency of $L_1$ average distribution

The result of the previous section shows that, given $n$ distributions $P^1$, $P^2$, ..., $P^n$ and aggregation matrix $W$, the barycenter of $WP^1$, $WP^2$, ..., $WP^n$ in $L_1$UniFrac space equivalent to their component-wise median. However, this does not guarantee a biologically meaningful average distribution with respect to $L_1$UniFrac. We provide an example of this below.

**Example 1.** *Consider* $P^1 = (0.3, 0, 0, 0.7)$*,* $P^2 = (0, 0.3, 0.2, 0.5)$*,* $P^3 = (0.4, 0.4, 0.2, 0)$ *and* $W = \begin{bmatrix} .9 & 0 & 0 & 0 \\ 0 & .9 & 0 & 0 \\ .1 & .1 & .1 & 0 \\ 1 & 1 & 1 & 1 \end{bmatrix}$.

We have $WP^1 = (0.27, 0, 0.03, 1)$, $WP^2 = (0, 0.27, 0.05, 1)$, $WP^3 = (0.36, 0.36, 0.1, 1)$ and the barycenter $WP^* = (0.27, 0.27, 0.05, 1)$. However, $W^{-1}P^* = (0.3, 0.3, -0.1, 0.5)$, which contains a negative entry, despite the fact that the entries sum up to 1. This means that the $W^{-1}P^*$ fails to uphold the biological meaning of representing an average microbial distribution.

This issue can be avoided when $L_2$UniFrac is used instead of $L_1$UniFrac.

##### S1.4.2 The advantages of $L_2$ average distribution

**Claim 3.** *The barycenter of a finite number of vectors in $L_2$ space is equivalent to component-wise mean of the vectors.*

*Proof.* Let $x^1$, $x^2$, ..., $x^n \in (\mathbb{R}^N, ||.||)$ be $n$ vectors in $L_2$ space, where $||.||$ is the $L_2$-norm. Then the barycenter $x^*$ satisfies

$$x^* = \arg\min_x \sum_{i=1}^{n} \sqrt{\sum_{j=1}^{N} (x_j^i - x_j)^2} \tag{8}$$

$$\Rightarrow x_j^* = \arg\min_x \sum_{i=1}^{n} (x_j^i - x_j)^2 \tag{9}$$

This implies $\sum_{i=1}^{n}(x_j^i - x_j)^2$ attains its minimum when $x_j = x_j^*$. Taking the derivative of $\sum_{i=1}^{n}(x_j^i - x_j^*)^2$, we have

$$\sum_{i=1}^{n}(x_j^i - x_j^*) = 0 \tag{10}$$

$$\sum_{i=1}^{n} x_j^i - n x_j^* = 0 \tag{11}$$

$$x_j^* = \frac{1}{n}\sum_{i=1}^{n} x_j^i. \tag{12}$$

$$\tag{13}$$

□

**Claim 4.** *Given $n$ probability vectors $x^1, x^2, ..., x^n$ in $\mathbb{R}^N$ satisfying $x_j^i \geq 0$ $\forall 1 \leq j \leq N$ and $\sum_{j=1}^{N} x_j^i = 1$ $\forall 1 \leq i \leq n$, and an invertible matrix $W \in \mathbb{R}^{\mathbb{N}\times\mathbb{N}}$, let $x^*$ denote the $L_2$-average vector obtained as described above by successively applying $W$, taking the average, and applying $W^{-1}$. Then $x^*$ is a probability vector. Namely, $x_j^* \geq 0$ $\forall 1 \leq j \leq N$ and $\sum_{j=1}^{N} x_j^* = 1$.*

*Proof.* Let $\overline{x}$ denote the mean vector of the $L_2$-aggregated vectors in the $L_2$UniFrac space after applying $W$. Namely,

$$\overline{x} = \frac{1}{n}\sum_{i=1}^{n} W x^i \tag{14}$$

$$\Rightarrow \overline{x}_j = \frac{1}{n}\sum_{i=1}^{n}(w_{j1}x_1^i + w_{j2}x_2^i + ... + w_{jN}x_N^i) \tag{15}$$

$$= \frac{1}{n}(w_{j1}\sum_{i=1}^{n} x_1^i + w_{j2}\sum_{i=1}^{n} x_2^i + ... + w_{jN}\sum_{i=1}^{n} x_N^i).$$

Since $W$ is invertible, there exists $W^{-1}$ with $k, l$-entry denoted by $w'_{kl}$ such that $\overline{W} := W^{-1}W$ satisfies

$$\overline{w}_{kl} = \sum_{j=1}^{N} w'_{kj} w_{jl} = \begin{cases} 1 & \text{if } k = l \\ 0 & \text{otherwise} \end{cases}. \tag{16}$$

By definition,

$$x^* = W^{-1}\overline{x} \tag{17}$$

$$\Rightarrow x_j^* = (w'_{j1}\ w'_{j2}\ ...\ w'_{jN}) \cdot \overline{x}$$

$$= \frac{1}{n}(w'_{j1}\ w'_{j2}\ ...\ w'_{jN}) \cdot \begin{bmatrix} \sum_{j=1}^{N}(w_{1j}\sum_{i=1}^{n} x_1^i) \\ \sum_{j=1}^{N}(w_{2j}\sum_{i=1}^{n} x_2^i) \\ \vdots \\ \sum_{j=1}^{N}(w_{Nj}\sum_{i=1}^{n} x_N^i) \end{bmatrix}$$

$$= \frac{1}{n}(\sum_{k=1}^{N} w'_{jk}\sum_{j=1}^{N} w_{1j}\sum_{i=1}^{n} x_k^i)$$

$$= \frac{1}{n}(\sum_{i=1}^{n} x_1^i \sum_{k=1}^{N} w'_{jk}w_{k1} + \sum_{i=1}^{n} x_2^i \sum_{k=1}^{N} w'_{jk}w_{k2} + ... + \sum_{i=1}^{n} x_N^i \sum_{k=1}^{N} w'_{jk}w_{kN})$$

$$= \frac{1}{n}\sum_{i=1}^{n} x_j^i \sum_{k=1}^{N} w'_{jk}w_{kj} \text{ by (16)}$$

$$= \frac{1}{n}\sum_{i=1}^{n} x_j^i. \tag{18}$$

Since $x_j^i \geq 0 \forall i, j$, $x_j^* \geq 0 \forall j$. Also, $\sum_{j=1}^{N} x_j^* = \frac{1}{n}\sum_{i=1}^{n}\sum_{j=1}^{N} x_j^i = \frac{1}{n}(n) = 1$. □

This guarantees that the barycenter obtained with respect to $L_2$UniFrac will always be a distribution. Next, we show that not only is this $x^*$ an distribution, it is also the barycenter of $x^1$ through $x^n$ in $(\mathbb{R}^N, d)$ where $d$ is the $L_2$UniFrac metric.

**Claim 5.** *Given probability distributions $x^1$, $x^2$, ..., $x^n$ in $\mathbb{R}^N$ and aggregation matrix $W$, let $y$ be the barycenter of $Wx^1$, $Wx^2$, ..., $Wx^n$ in $L_2$ UniFrac space. Then $W^{-1}y$ is the barycenter of $x^1$, $x^2$, ..., $x^n$ in the distribution space under the $L_2$ UniFrac metric.*

*Proof.* Let $x^*$ be the barycenter of $x^1$, $x^2$, ..., $x^2$ under $L_2$UniFrac.Then

$$x^* = \arg\min_x \sum_{i=1}^{n} ||W(x^i - x)|| \tag{19}$$

$$= \arg\min_x \sum_{i=1}^{n} ||W(x^i - x)||^2 \tag{20}$$

$$= \arg\min_x \sum_{i=1}^{n} \sum_{j=1}^{N} (\sum_{k=1}^{N} w_{jk}(x_k^i - x_k))^2 \tag{21}$$

$$= \arg\min_x \sum_{i=1}^{n} (\sum_{k=1}^{N} w_{jk}(x_k^i - x_k))^2 \quad \forall 1 \leq j \leq N \tag{22}$$

$$\Rightarrow \sum_{i=1}^{n} \sum_{k=1}^{N} w_{jk}(x_k^i - x_k^*) = 0 \tag{23}$$

$$\sum_{k=1}^{N} w_{jk}(\sum_{i=1}^{n} x_k^i - nx_k^*) = 0. \tag{24}$$

where the derivative is taken in line (23) and is known to be zero as it's at a minimum. Since $W$ is an arbitrary aggregation matrix, it has to be the case that $\sum_{i=1}^{n} x_k^i = nx_k^*$. Which in turn implies that $x_k^* = \frac{1}{n}\sum_{i=1}^{n} x_k^i$ for all $k$. By claim 4, $x^* = W^{-1}y$. □

Moreover, (18) shows that the $L_2$-average distribution obtained in this way is equivalent to taking the $L_2$ mean of the original distribution component-wise. This nice property is again absent in $L_1$UniFrac, as can be seen in this simple counter-example of $P^1 = (1, 0, 0)$, $P^2 = (0.17, 0.33, 0.5)$ and $P^3 = (0.33, 0.5, 0.17)$, of which the component-wise median equates to $(0.33, 0.33, 0.17)$, which does not have components summing up to 1.

#### S1.5 Barycenter with respect to $L_2$UniFrac metric

In this paper, we introduced the use of $L_2$UniFrac metric in place of the 1-Wasserstein-equivalent $L_1$UniFrac, and discussed its biological advantage of being able to yield the average distribution. In this section, we will discuss the significance of our results in a broader perspective.

Despite its popularity in measuring the dissimilarity between probability distributions, the Wasserstein distance is difficult to compute in general, having a cubic time complexity with respect to the number of supports [12]. Various attempts have been made to reduce the computational cost, either through approximation [10, 7], or projection to lower dimensional space [8], or looking for special cases where the computation can be simplified, such as the case of tree-Wasserstein distance [3]. We give a brief description of tree-Wasserstein distance.

**Definition 2** (Tree metric)**.** Given a finite set $\Omega$, a metric $d : \Omega \times \Omega \to \mathbb{R}$ is a tree metric if there exists a tree $T$ with non-negative edge lengths such that any element in $\Omega$ can be represented as a node on $T$ and for any $x, y \in \Omega$, $d(x, y)$ can be represented as a (unique) path between nodes $x$ and $y$ on $T$. [3]

A tree-Wasserstein metric is simply a measurement of the optimal transport with a the underlying metric being a tree metric. Under these definitions, it can be noticed that the ($L_1$) UniFrac metric, being equivalent to the 1-Wasserstein distance, is specifically a tree-Wasserstein metric with the phylogenetic tree being the underlying tree structure. The tree-Wasserstein metric is a special case of the Wasserstein metric due to the fact that it has a closed a form and can thus be computed in linear time [5, 3, 12].

Another important question related to the Wasserstein metric is finding the barycenter in a Wasserstein metric space. The Wasserstein barycenter has been a widely studied topic and finds its application in diverse fields such as natural language processing [14], model ensembling [1], and image processing [11, 9]. As can be expected, the computation of Wasserstein barycenter is closely related to the computation of the Wasserstein metric itself. Given the relative ease of computing tree-Wasserstein metrics, there has been a fair amount of effort invested in studying tree-Wasserstein barycenters. Though various methods have been proposed to speed up the computation of tree-Wasserstein barycenters, finding the exact barycenters remain a hard problem [12]. Most of the proposed methods still involve complicated algorithms and constrains, and are approximations in nature [12, 4]. Similarly for the case of $L_1$UniFrac, efficient computation of the exact barycenter remains an open problem, as demonstrated in Section S1.4.1. However, by simply changing the metric from $L_1$UniFrac to $L_2$UniFrac, we are able to obtain the exact barycenter with respect to this metric by simply taking the component-wise mean.

### S2 Supplementary figure

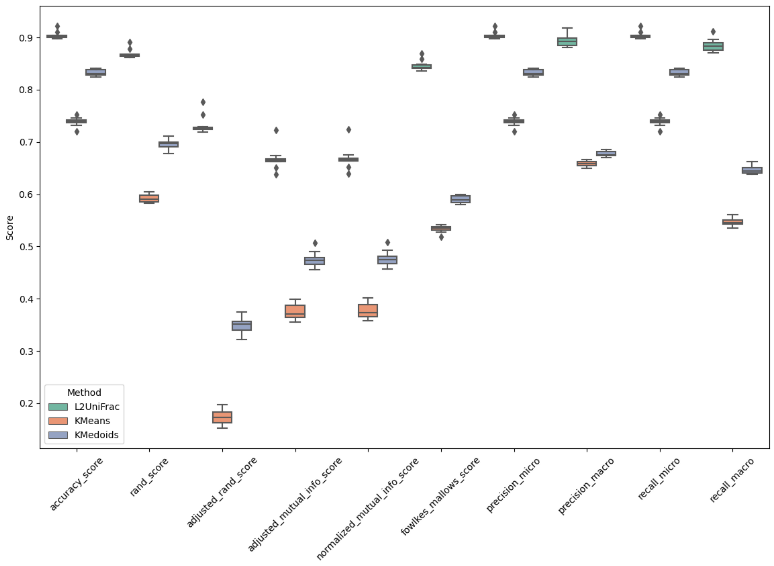


Figure S1: Other scores evaluating classification performance. Higher score indicates a better performance in all cases.
